# In-vitro activity of Cefiderocol against clinical isolates of meropenem-resistant *Klebsiella pneumoniae* from India

**DOI:** 10.1101/2022.09.16.508352

**Authors:** Kalyani Borde, M A Kareem, Ratna Mani Sharma, S Manick Dass, Vedantham Ravi, Dilip Mathai

**Affiliations:** Department of Microbiology, Apollo Institute of Medical Sciences and Research, Hyderabad; Department of Internal Medicine and Adult Infectious Diseases, Apollo Institute of Medical Sciences and Research, Hyderabad

**Keywords:** Cefiderocol, Carbapenem resistance, *Klebsiella pneumoniae*

## Abstract

**Background:** Cefiderocol (FDC), a novel siderophore drug, is active against gram negative bacteria producing carbapenemases, including metallo-beta-lactamases.

**Objective:** To compare the in-vitro activity of FDC with ceftazidime-avibactam (CZA), CZA/ aztreonam (AT) combination and colistin (CST), in clinical isolates of meropenem-resistant (MER-R) *Klebsiella pneumoniae*.

**Methods:** From 2052 clinical specimens submitted for culture testing, 245 K. pneumoniae isolates were recovered within a six month period in 2021. 103 non-duplicate, non-outbreak, MER-R (MIC >4 μg/ml) strains were included in the study. Identification and susceptibility was performed using VITEK-2 (bioMérieux). 10 meropenem-susceptible isolates served as controls. For FDC, BMD was performed after in-house standardisation. Disc diffusion (Liofilchem, Italy) and broth microdilution (BMD; ComASP, STC, Liofilchem, Italy) were used for susceptibility testing of CZA and CST respectively. Synergy testing for CZA and aztreonam (AT) was performed using disk approximation method. CLSI breakpoints were used for interpretation of results.

**Results:** For FDC, MIC_50_ and MIC_90_ was 2 μg/ml and 8 μg/ml, respectively. 80% isolates were susceptible to FDC. 26.2% isolates were susceptible to CZA, synergy testing with CZA/ AT was positive for 74 (72%) of the isolates. 89.3% had intermediate susceptibility to CST. Nine (8.7%) were susceptible only to FDC.

**Conclusion:** FDC is active in-vitro against MER-R *K. pneumoniae* > CZA/AT> CZA > CST, as observed in this study, applying CLSI criteria. Clinico-microbiological studies should be performed for assessing clinical efficacy of this novel drug in this region with high prevalence of carbapenem resistance among gram-negative organisms.

## Introduction

Cefiderocol (FDC) is a novel siderophore, with an ingenious description of its mechanism of action as ‘Trojan horse’. The drug enters the bacterial cell via active iron transporters, which helps overcome beta-lactamase - mediated resistance in gram negative organisms (1). The Infectious Disease Society of America (IDSA) and European Society for Clinical Microbiologists and Infectious Diseases (ESCMID) have recommended this drug for treating carbapenem-resistant Enterobacterales (CRE) (2, 3). Various large-scale trials have demonstrated high-level in-vitro activity of this drug against carbapenem-resistant organisms (4, 5). However, the data from countries such as India, with high burden of carbapenem resistance, particularly metallo-beta-lactamase (MBL) - mediated resistance, is lacking. According to the Indian national surveillance data, carbapenem resistance is high among clinical isolates of *Klebsiella pneumoniae* (6). The aim of the study was to evaluate in-vitro activity of FDC, and compare it with currently available therapy options such as ceftazidime-avibactam (CZA) and colistin (CST).

## Material and method

### Place of study

The study was carried out prospectively at the microbiology laboratories attached to a teaching hospital and a tertiary care centre in southern India, during 2021. The study was approved by the institutional research and ethical committee (AIMSR/ IRB/ 2020/ 11/ B/ 7).

### Isolate selection

From the clinical samples submitted for bacterial cultures, 113 non-repetitive isolates of *K. pneumoniae* as identified by VITEK-2 were selected. Meropenem susceptibility was performed using VITEK-2 GN cards. Other antibiotics tested with VITEK-2 were amikacin, ampicillin, amoxicillin-clavulanic acid, cefuroxime, ceftriaxone, cefepime, ciprofloxacin, cotrimoxazole, gentamicin, imipenem, nitrofurantoin, piperacillin-tazobactam, tigecycline and CST. Isolates were stored at -80°C in glycerol broth, until further testing.

### MIC testing

FDC drug in pure form (research-use-only) was obtained from Shionogi & Co. Ltd. Dilutions were prepared in round bottom microtiter plates, ranging from 0.03 μg/ml to 32 μg/ml. This range was selected to cover the MIC ranges of quality control strains at the lower end. The plates were stored at -80°C. When sufficient isolates were accumulated, plates were thawed and inoculated. For preparing inoculum, research-use-only iron-deficient, cation adjusted Mueller-Hinton broth (ID-CAMHB; Thermo Fisher, The USA) was used. BMD plates were incubated at 35 ± 2 °C for 16-20 hours. Results were interpreted as the first well where visible bacterial growth was inhibited (Image 1). In case of trailing growth (multiple wells of faint growth as compared to the growth control well), the well with 80% inhibition of growth was taken as the MIC. All 113 isolates were tested for FDC MIC using BMD. Colistin MIC testing was performed for all 113 isolates using commercial lyophilized broth microdilution plates (Liofilchem®, Italy) for the range 0.25 μg/ml to 16 μg/ml. CLSI criteria were used for interpretation (7)

### Disk diffusion testing

FDC (30 μg), CZA (30/20 μg) and aztreonam (30 μg) disks were tested as per Kirby-Bauer disk diffusion method on Meuller-Hinton agar (MHA). Zone diameters were read after incubation at 35 ± 2 °C for 16-18 hours. FDC disk diffusion was performed for 52 isolates as the disks were received at a later date due to COVID-related delay in shipments. Synergy between CZA and AT was tested for all 103 isolates by observing zone distortion (Image 2) between the disks of CZA and AT placed 20 mm apart.

### Quality control

For MIC as well as disk diffusion testing of FDC, American Type Culture Collection (ATCC^®^) strains of *E. coli* 25922 and *P. aeruginosa* 27853 were used. For CST susceptibility testing, National Collection of Type Cultures (NCTC) *E. coli* 13846 was used.

### Agreement analysis

Taking BMD as the gold standard, categorical agreement (CA) with the disk diffusion was calculated. Very major errors (VME), major errors (ME) and minor errors (mE) were also calculated.

## Results

Of the 113 clinical isolates of *K. pneumoniae*, 37%, 22%, 20%, 14% and 7% of the isolates were from hospitalised patients with blood, urine, respiratory tract, pus, and from other sites, respectively.

### FDC susceptibility testing

FDC MICs for meropenem-susceptible isolates (n=10) ranged from ≤ 0.03 μg/ml to 2 μg/ml. For meropenem-resistant isolates (n=103), MIC_50_ and MIC_90_ were 2 μg/ml and 8 μg/ml, respectively. Of the 103, 83 (80.6%) were susceptible to FDC (MIC ≤ 4 μg/ml), 15 (14.5%) showed intermediate susceptibility (MIC = 8 μg/ml) and 5 (4.8%) showed resistance (MIC ≥ 16 μg/ml), according to CLSI breakpoints (Figure 1).

**Figure 1:**
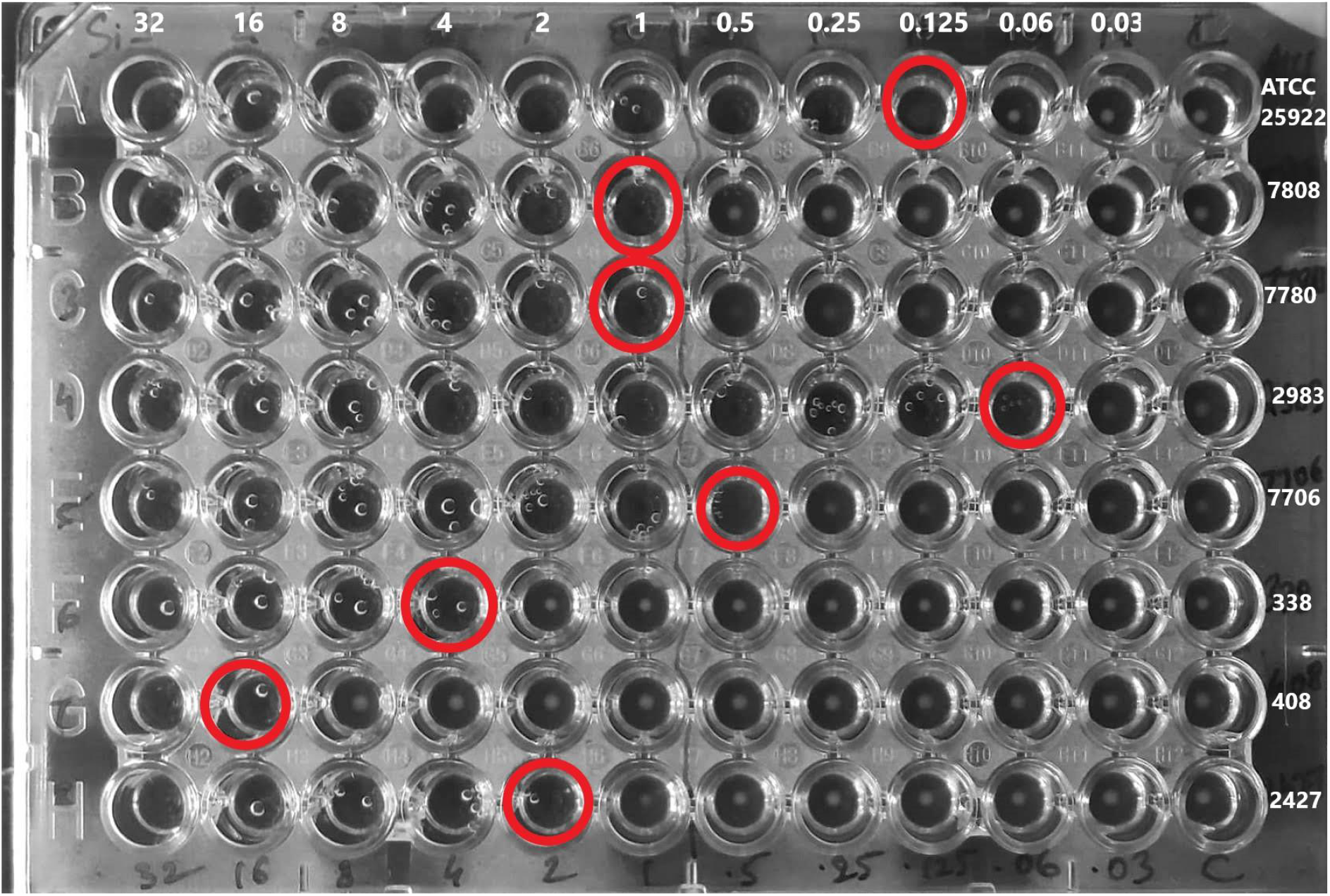
Reading of cefiderocol broth microdilution testing.

**Figure 2:**
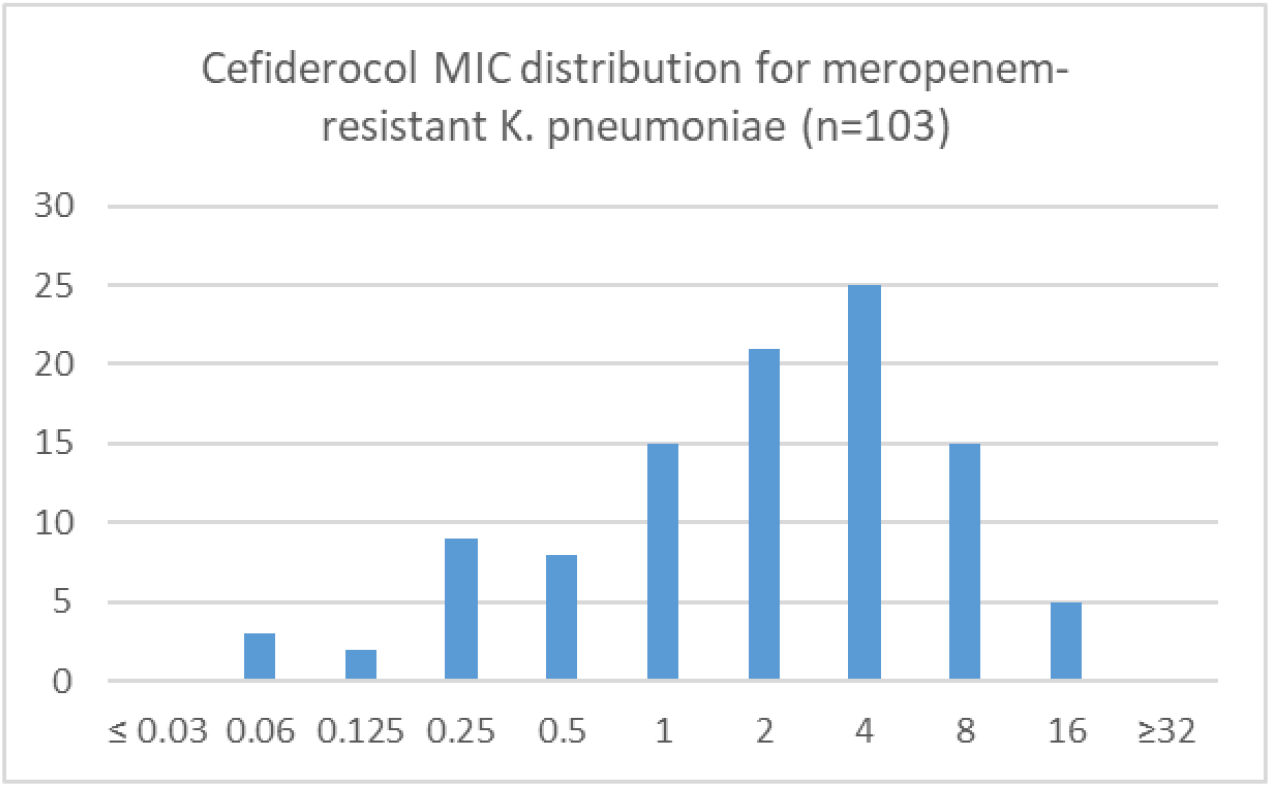
Distribution of minimum inhibitory concentrations of meropenem-resistant *Klebsiella pneumoniae* isolates included in the study.

FDC disk diffusion was performed for 52 of these 103 isolates (Table 1). With the CLSI breakpoints, categorical agreement with BMD was 90%, 50% and 100% for susceptible, intermediate and resistant categories, respectively. Major errors (ME) occurred in 1.9% and minor errors (mE) in 9.6%. There were no very major errors (VME).

**Table 1:** Relative distribution of zone diameters and minimum inhibitory concentrations of meropenem-resistant Klebsiella pneumoniae strains included in the study.

| Zones (mm)→<br>MIC (µg/ml)<br>↓ | 7 | 8 | 9 | 10 | 12 | 14 | 15 | 16 | 17 | 18 | 19 | 20 | 22 | 23 | 24 | 25 | 26 | 27 | 29 | 30 | 32 | 33 |
| --- | --- | --- | --- | --- | --- | --- | --- | --- | --- | --- | --- | --- | --- | --- | --- | --- | --- | --- | --- | --- | --- | --- |
| 0.03 |  |  |  |  |  |  |  |  |  |  |  |  |  |  |  |  |  |  |  |  |  |  |
| 0.06 |  |  |  |  |  |  |  |  |  |  |  |  |  |  | 1 | 1 |  |  |  |  |  |  |
| 0.125 |  |  |  |  |  |  |  |  |  |  | 1 |  |  |  | 1 |  |  |  |  |  |  |  |
| 0.25 |  |  |  |  |  |  |  |  |  |  |  |  |  |  | 1 |  | 1 |  |  |  | 1 |  |
| 0.5 |  |  |  |  |  |  |  |  |  |  |  |  |  |  |  |  |  | 1 | 2 |  |  |  |
| 1 |  |  |  |  |  |  |  |  | 1 |  |  |  |  | 2 |  |  | 3 | 1 | 1 |  | 1 |  |
| 2 |  |  |  |  | 1 | 1 | 1 |  |  | 1 |  |  | 3 |  |  | 1 |  | 1 |  |  |  | 1 |
| 4 | 1 |  |  |  |  |  |  |  |  | 1 |  | 1 | 1 | 2 |  | 3 | 2 |  |  | 1 |  |  |
| 8 |  |  |  |  | 1 | 1 | 2 | 1 |  | 1 | 1 |  |  |  |  | 1 |  |  |  |  |  |  |
| 16 |  | 1 | 1 | 1 |  |  |  |  |  |  |  |  |  |  |  |  |  |  |  |  |  |  |
| 32 |  |  |  |  |  |  |  |  |  |  |  |  |  |  |  |  |  |  |  |  |  |  |

Note: Zone diameters in millimeters are in the columns and minimum inhibitory concentrations (MIC) in µg/ml are in the rows. Red, yellow and green colors indicate resistant, intermediate and susceptible zone diameters or concentrations, respectively. (CLSI criteria, 2022).

Resistance was found in 43% when EUCAST breakpoints for MIC testing were applied (8). Similarly, categorical agreement with disk diffusion was 79% and 56% for susceptible and resistant categories, respectively. With EUCAST breakpoints, VME occurred in 19.2% and ME occurred 11.5%. (Table 2, 3).

**Table 2:** Categorical agreement between broth microdilution and disk diffusion for cefiderocol according to CLSI and EUCAST breakpoints.

| **CLSI** | Susceptible | Intermediate | Resistant |
| --- | --- | --- | --- |
| BMD | 41 | 8 | 3 |
| DD | 37 | 4 | 1 |
| CA | 90% | 50% | 33% |
| **EUCAST** |  |  |  |
| BMD | 29 | Not applicable | 23 |
| DD | 23 |  | 13 |
| CA | 79% |  | 56% |

BMD: Broth microdilution
DD: Disk diffusion
CA: Categorical agreement

**Table 3:** Error rates for disk diffusion for CLSI and EUCAST breakpoints.

|  | **CLSI** |  | **EUCAST** |  |
| --- | --- | --- | --- | --- |
| **VME** | 0 | 0.00% | 10 | 19.20% |
| **ME** | 1 | 1.90% | 6 | 11.50% |
| **mE** | 5 | 9.60% | Not applicable |  |

VME: Very major error - when resistant (R) is categorised as susceptible (S)
ME: Major error - when susceptible (S) is categorised as resistant (R)
mE: Minor error - when susceptible (S) or resistant (R) is categorised as intermediate susceptible (I).

Nine (8.7%) isolates were susceptible only to FDC, and resistant to all other antibiotics tested.

### CST broth microdilution testing

Of the 103 meropenem resistant K. pneumoniae, 92 (89%) had intermediate susceptibility to CST (MIC ≤ 2 μg/ml) and 11 (11%) were resistant (≥ 4 μg/ml). For three of these 11 resistant isolates, discrepancies were observed with VITEK-2 MICs, two isolates were categorised as intermediate (MIC ≤ 2 μg/ml) by VITEK-2.

Four (3.8%) isolates were intermediate susceptible only to colistin, and resistant to all other antibiotics tested.

### CZA susceptibility testing

Of the 103 meropenem-resistant K. pneumoniae, 27 (26.2%) were susceptible, 1 (1%) had intermediate susceptibility and 75 (72.8%) were resistant to CZA.

### Synergy testing between CZA and AT

Synergy testing was positive for 74 (72%) of the isolates. Out of these 74, 59 (80%) isolates were resistant when tested individually for both CZA and AT. Synergy was present in 14 out of 20 (70%) isolates which were non-susceptible (I, R) to FDC. Among the five isolates resistant (R) to FDC, only one showed synergy between CZA and AT.

## Discussion

The national AMR surveillance network in India has shown the susceptibility of only 47% against meropenem for K. pneumoniae (6). This seriously limits the treatment options, necessitating use of last resort antibiotics.

FDC is an FDA-approved, parenteral, siderophore cephalosporin antibiotic, which acts by interfering with iron uptake by the microorganisms (9). The structure combines a catechol moiety with a cephalosporin core, giving it enhanced stability against many β-lactamases including MBL (10). It is recommended for treatment of carbapenem-resistant Enterobacterales in regions with high prevalence of MBL. Susceptibility testing of FDC by BMD requires use of iron-deficient media, as iron interferes with the activity of this drug. This can be technically challenging, lowering the reliability and reproducibility of in-house testing. However, disk diffusion is performed without any additional requirements and is shown to be reliable (11).

Colistin belongs to the polymyxin class of antibiotics. The target serum concentrations are as high as 2 ug/ml for it to be effective, which are not achieved in more than 50% of the patients and result in nephrotoxicity (12). The susceptibility testing of colistin is also fraught with issues such as low reproducibility and heteroresistance. Disk diffusion, gradient testing or semi-automated commercial testing panels (e.g. VITEK-2) are not recommended for testing this drug. The only recommended method is broth microdilution, which is not widely available, especially in resource-limited settings. Khurana et. al. found 10% VMEs with VITEK-2 (13). In our study, we also observed discrepancies between BMD and VITEK-2 for colistin susceptibility, with lower VMEs at 1.9%. CLSI has also removed the category ‘susceptible’ for this drug, keeping only ‘intermediate’ and ‘resistant’ categories for reporting (7). In our study, 89% of the isolates had intermediate susceptibility to colistin. This is in line with the findings of Amladi et al and Manohar et al (14, 15).

Ceftazidime-avibactam is a novel drug which is shown to be effective against carbapenem-resistant Enterobacterales, except those producing MBLs (16). Since India has reported a high prevalence of MBLs (17), this drug alone is not very effective but can be used in combination with aztreonam (2). Low susceptibility was observed in our study, with only 26% isolates being susceptible to CZA. Susceptibility testing is easily performed with routine disk diffusion. However, there are no guidelines for synergy testing between CZA and AT. A study by Sahu et. al. (18) found more than 80% of the isolates showing synergy for CZA/ AT and 96% of these harboured *bla*_NDM-1_. In comparison, synergy testing was positive in 72% of our isolates.

Our study shows less susceptibility (MIC_90_ of 8 μg/ml) than the SIDERO-WT study, (MIC_90_ of 4 μg/ml) for carbapenem-nonsusceptible Enterobacterales (5). However, the study represented isolates majorly obtained from Europe and North America, with only 8% isolates originating from Asia. Similarly, cefiderocol susceptibility performed on isolates collected during the SENTRY study showed MIC_90_ of 4 μg/ml for CRE, with CRE constituting only 2.1% of the study isolates(19). In another study from China, among 105 carbapenem-resistant K. pneumoniae, 100% were susceptible to FDC. However, only 8 isolates harboured *bla*_NDM-1_, remaining harboured *bla*_KPC_. Isolates harbouring *bla*_NDM-1_ were found to have higher MICs for FDC in this study (20). Co-production of multiple carbapenemases could contribute to resistance as discussed by Kohira et. al. (21). Indian isolates often harbour more than one carbapenemase gene (17, 22). This might have contributed to lower susceptibility in our study than that in the western literature. Furthermore, resistance to CZA might contribute to reduced susceptibility to FDC, as noted by Bianco et al (23).

In our study, there was a wide variation between susceptibility rates when applying CLSI and EUCAST breakpoints for FDC. 80% and 56% of the isolates were susceptible to FDC, when CLSI and EUCAST breakpoints were used respectively. These findings are in line with Morris et. al. (24). Disk diffusion is a convenient alternative to MIC testing for FDC, since BMD testing for the same is fraught with many technical challenges (25). Matuschek et al have validated disk diffusion as a reliable and robust alternative to BMD (11). However, in our study, categorical agreement varied widely between CLSI (CA = 80%) and EUCAST (CA = 69%) breakpoints. Error rates also varied widely with CLSI breakpoints giving less error rates as compared to the EUCAST criteria. This variation is also observed by Morris et al and Bonnin et al (24, 26). Hence, it can be argued that the breakpoints need to be investigated further for harmonisation, keeping in view the clinical response and PK-PD parameters, especially in areas with a high burden of carbapenem resistance.

Among other treatment options for MER-R K. pneumoniae, tigecycline was not compared in the study, as more than 70% of our isolates were from blood, respiratory system or urine; all these are the sites where tigecycline is not effective. Fosfomycin was not compared as only 20% isolates were recovered from urine. Molecular characterization and disk diffusion of all the study isolates could not be performed, which are the limitations of the study.

## Conclusion

FDC exhibited reasonable in-vitro activity against meropenem-resistant *K. pneumoniae* in our study isolates. Larger clinico-microbiological studies should be performed for assessing clinical efficacy of this antibiotic in this region with high prevalence of carbapenem resistance among gram-negative organisms.

## Acknowledgement

Authors received the drug FDC in pure form for research-use-only from Shionogi co. Authors received ID-CAMHB from ThermoFisher for research-use-only. Neither had any bearing upon the study design or findings.

Authors acknowledge help by Dr. Gunnar Kahlmeter, Dr. Erika Matuschek (EUCAST Development Lab, Vaxjo, Sweden) and Dr. Mandy Wootton (Cardiff, Wales), offered via online correspondence, for troubleshooting the broth microdilution testing of FDC.

## Tables and Figures

**Image 2:**
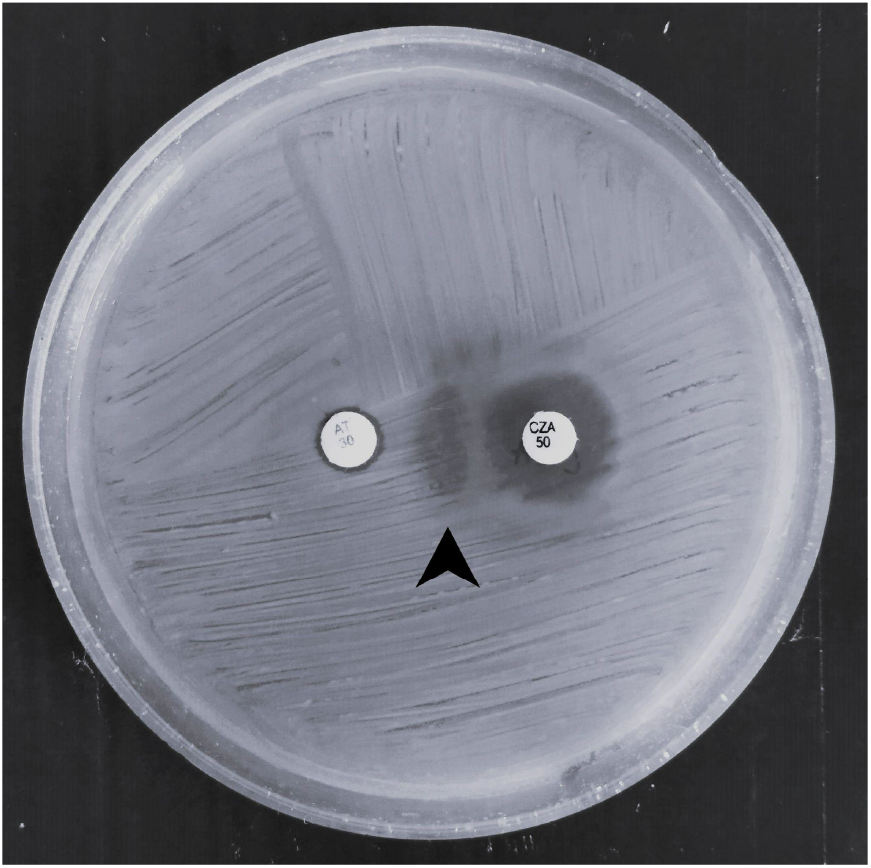
Synergy testing between aztreonam and ceftazidime-avibactam by disk diffusion method. Note: Arrowhead - Zone distortion showing synergy between aztreonam and ceftazidime-avibactam.

